# Metabolic defects cause hyperactive mitochondria and Parkinson’s disease-like traits

**DOI:** 10.1101/2020.02.20.958322

**Authors:** Danielle E. Mor, Salman Sohrabi, Rachel Kaletsky, Will Keyes, Vrinda Kalia, Gary W. Miller, Coleen T. Murphy

## Abstract

Metabolic dysfunction is a facet of many age-related neurodegenerative diseases, yet its role in disease etiology remains poorly understood^1^. We recently discovered a potential causal link between the branched-chain amino acid transferase, *BCAT-1*, and the neurodegenerative movement disorder, Parkinson’s disease (PD)^2^. Knockdown of *C. elegans bcat-1* recapitulates PD-like features, including progressive motor deficits and neurodegeneration with age^2^. Using transcriptomic, metabolomic, and imaging approaches, we show that *bcat-1* knockdown increases mitochondrial activity and induces oxidative damage in neurons through mTOR-independent mechanisms. We recently developed a high-throughput screening platform to identify drugs that may be repurposed for PD, and found that metformin, the leading type 2 diabetes medication, significantly improves motor function in *bcat-1(RNAi)* worms^3^. Late-in-life metformin treatment restores normal mitochondrial activity levels and protects against *bcat-1-*associated neurodegeneration. Our results suggest that PD may originate as a metabolic disorder, and highlight metformin as a promising new drug candidate for PD treatment.

---

Parkinson’s disease (PD) is a devastating neurodegenerative movement disorder, and its prevalence is predicted to significantly increase as the population ages^4,5^. PD is characterized by the progressive loss of dopaminergic neurons in the substantia nigra, and the formation of pathological inclusions containing aggregated α-synuclein protein^6,7^. The mechanisms that induce neurodegeneration in PD are still poorly understood, and greater than 90% of cases have no known cause^8^.

In an effort to identify novel PD genes, we recently developed *diseaseQUEST*, a method that combines human genome-wide association studies with tissue-specific functional genetic networks and high-throughput behavioral screening in *C. elegans*^2^. Using this computational-experimental framework, we discovered a novel link between branched-chain amino acid transferase 1 (*BCAT-1*), which catalyzes the first step in branched-chain amino acid (BCAA) catabolism^9^, and PD. We found that *BCAT-1* expression is normally high in PD-susceptible regions of the healthy human brain, and its expression is reduced in the substantia nigra of sporadic PD patients^2^. While these findings suggest a correlation between defective BCAA metabolism and PD, animal models are required to determine causality and underlying mechanisms.

The nematode *C. elegans* offers a unique system in which to study age-related neurological disease, due to its short lifespan and highly conserved nervous system signaling that gives rise to a set of complex behaviors. Moreover, the worm is highly amenable to high-throughput screening approaches, allowing rapid testing of potential disease treatment options^10^. Previously, we showed that RNAi-mediated reduction of neuronal *bcat-1* in *C. elegans* causes progressive, age-dependent motor dysfunction and accelerates dopamine neuron degeneration in worms expressing human α-synuclein^2^. Here, we use tissue-specific transcriptomics, high-resolution metabolomics, and imaging in aged worms, revealing that *bcat-1* knockdown - surprisingly - increases neuronal mitochondrial activity. The type 2 diabetes medication metformin restores normal mitochondrial activity levels, reduces neurodegeneration, and significantly improves motor function, even with late-in-life administration. Our work underscores the possibility that at least some forms of PD may primarily be a metabolic disorder, and offers metformin as a new potential treatment option for PD.

## Neuronal transcriptional analysis reveals upregulation of bioenergetic pathways

To model the reduction of *BCAT-1* that was observed in PD patient substantia nigra^2^, we performed adult-specific RNAi-mediated knockdown of *bcat-1* in *C. elegans* to avoid embryonic/larval lethality^11^. Consistent with our previous findings^2^, *bcat-1* knockdown exclusively in the nervous system (*sid-1;Punc-119::sid-1*) caused severe, spasm-like ‘curling’ motor dysfunction in aged (day 8 adult) animals (Fig. 1a). By this timepoint, dopamine neurons are almost entirely lost in neuronal RNAi-sensitive worms expressing dopaminergic-specific α-synuclein (Extended Data Fig. 1a); however, examination at the earlier timepoint of day 6 revealed that *bcat-1* knockdown exacerbated the loss of cell bodies and the fragmentation and blebbing of neurites (Fig. 1b). Dopaminergic degeneration is associated with loss of *C. elegans’* basal slowing response, in which worms normally slow down in the presence of a bacterial food source^12^. Consistent with the observed structural damage (Fig. 1b), *bcat-1* knockdown worsened basal slowing performance (Extended Data Fig. 1b).

**Figure 1:**
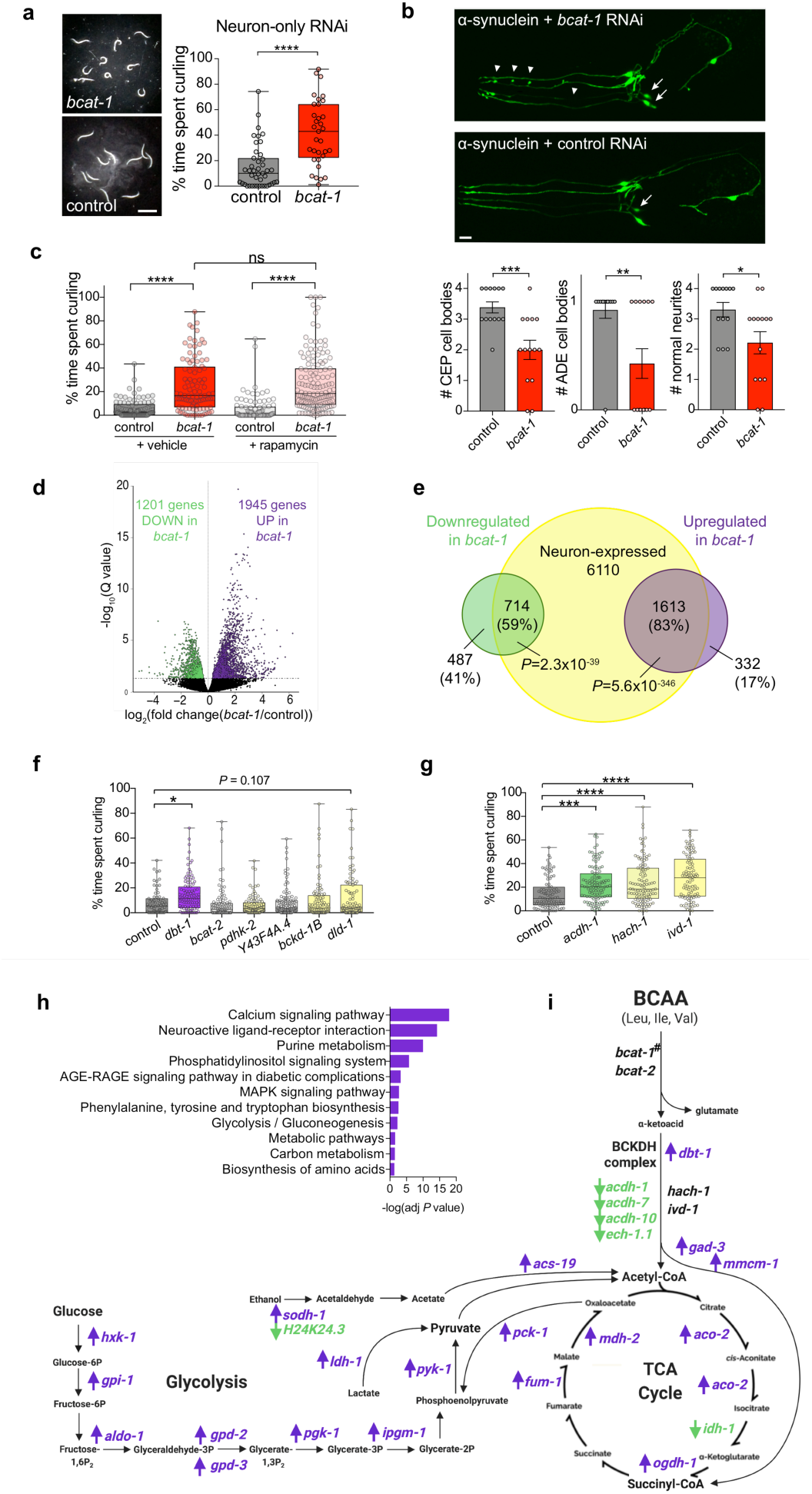
*bcat-1* knockdown upregulates bioenergetic pathway genes in neurons. **a**, Adult- and neuron-specific RNAi-mediated *bcat-1* knockdown results in severe spasm-like ‘curling’ motor dysfunction in aged (day 8 adult) animals. Scale bar, 1mm. n=43 for control, 33 for *bcat-1(RNAi)*. Two-tailed *t*-test **b**, In neuronal RNAi-sensitive worms expressing α - synuclein in dopaminergic neurons, *bcat-1* knockdown accelerates degeneration of CEP and ADE dopaminergic cell bodies (arrows) and neurites (arrowheads) on day 6. Scale bar, 10μm. n=13 for contjrol, 14 for *bcat-1 (RNAi).* Two-tailed *t*-test. Data are mean±s.e.m. **c**, Curling is unaffected by rapamycin treatment in neuronal RNAi-sensitive worms with *bcat-1* knockdown on day 8. ns, not significant. n=92 for control vehicle, 97 for *bcat-1* vehicle, 87 for control rapamycin, 139 for *bcat-1* rapamycin. Two-way ANOVA with Tukey’s post-hoc. **d-e**, Neurons were isolated on day 5 from neuronal RNAi-sensitive worms with adult-specific *bcat-1* knockdown and RNA-sequenced. n=5 independent collections/group. Volcano plot **(d)** showing upregulated (purple) and downregulated (green) genes in *bcat-1(RNAi)* neurons (FDR<0.05). Venn diagram **(e)** showing a majority of differentially-expressed genes are expressed in neurons. *P* values: hypergeometric distributions, **f-g**, Adult-specific knockdown of BCAA pathway components in neuronal RNAi-sensitive worms showed disruption of **dbt-1** *(f), acdh-1, hach-1*, or *ivd-1* **(g)** causes curling on day 8. n=70 and 99 for control in **(f)** and **(g)**, respectively, 100 for *dbt-1*, 95 for *bcat-2*, 91 for pdhk-2, 111 for *Y43F4A.4*, 85 for *bckd-1B*, 90 for *dld-1*, 111 for *acdh-1*, 107 for *hach-1*, 96 for *ivd-1*. One-way ANOVA with Dunnett’s post-hoc. **h**, KEGG analysis of genes upregulated in *bcat-1(RNAi)* neurons, **i**, BCAA, glycolysis, and TCA cycle pathway schematic with genes upregulated (purple) and downregulated (green) in *bcat-1(RNAi)* neurons, ^#^*bcat-1* expression could not be measured due to RNA-sequencing of the *bcat-1* RNAi. \**P*<0.05, \*\**P*<0.01, \*\*\**P*<0.001, 0.0001. Box-plots show minimum, 25th percentile, median, 75th percentile, maximum.

To understand how *bcat-1* reduction causes neurodegeneration and motor deficits, we first investigated the mTOR pathway, which was previously implicated in *bcat-1* regulation of lifespan^13^. However, we found that this pathway does not interact with *bcat-1* to regulate PD-related phenotypes, since neither mTOR/*let-363* RNAi nor rapamycin treatment had any effect on curling behavior (Fig. 1c, Extended Data Fig. 2).

Therefore, we decided to take an unbiased and tissue-specific approach to uncover the mechanisms of *bcat-1* neurotoxicity, using our method for the isolation and RNA sequencing of adult *C. elegans* tissues^14^. Neuronal RNAi-sensitive *C. elegans* were treated with *bcat-1* or control RNAi, and neurons were collected on day 5 of adulthood, a timepoint that precedes severe motor dysfunction (Fig. 1a), in order to shed light on mechanisms that drive disease rather than reporting transcriptional changes from dying neurons. *bcat-1(RNAi)* and control RNAi-treated samples were distinct (Extended Data Fig. 3a), and the majority of the differentially-expressed (FDR<0.05) genes (Fig. 1d, Extended Data Table 1) were previously identified as neuronally-expressed in adult wild-type *C. elegans*^15^ (Fig. 1e). Gene Ontology analysis also revealed that genes upregulated in *bcat-1(RNAi)* neurons were largely neuronal, including terms such as *chemical synaptic transmission, synapse organization*, and *regulation of neurotransmitter levels* (Extended Data Fig. 3b,c).

Several genes involved in BCAA metabolism were differentially expressed in response to *bcat-1* knockdown (Fig. 1i, Extended Data Table 1). To test if disrupting these BCAA pathway components produces motor dysfunction similar to *bcat-1(RNAi)*, neuronal RNAi-sensitive worms were fed RNAi as adults and tested for curling on day 8. The acyl-CoA dehydrogenase *acdh-1* was the most downregulated neuronal gene (log_2_(fold-change)=-2.44, p-adj=2.87×10^-5^), while *dbt-1*, which encodes the catalytic core of the branched chain α-ketoacid dehydrogenase complex (BCKDHC), was increased in *bcat-1(RNAi)* neurons (log_2_(fold-change)=0.55, p-adj=0.04). Disruption of either *acdh-1* or *dbt-1* was sufficient to induce curling (Fig. 1f,g). Since *acdh-1* functions in valine and isoleucine metabolism but not leucine^16^, we also tested *hach-1*, a hydrolase specific to valine, and *ivd-1*, a dehydrogenase specific to leucine, and in each case, knockdown was sufficient to produce curling (Fig. 1g), indicating that defects in the metabolism of any of the three BCAAs can produce motor dysfunction. Therefore, normal BCAA metabolism appears to be critical to neuronal maintenance with age.

KEGG functional analysis of genes upregulated in *bcat-1(RNAi)* neurons suggested enrichment of several metabolic pathways, including *glycolysis/gluconeogenesis, carbon metabolism, metabolic pathways*, and *purine metabolism* (Fig. 1h, Extended Data Fig. 3d). We further examined carbon metabolism, in particular glycolysis and the TCA cycle, since BCAA metabolism is closely linked with these bioenergetic pathways^9^ (Fig. 1i) and BCAT-1 localizes to mitochondria in *C. elegans*^17^. The majority (19/25) of the differentially-expressed genes in these pathways were upregulated in *bcat-1(RNAi)* neurons (Fig. 1i), suggesting that *bcat-1* knockdown may promote a coordinated transcriptional response to alterations in BCAA metabolism.

## Metabolomics shows TCA cycle depletion while mitochondrial activity is increased

To better understand the effects of *bcat-1* knockdown on carbon metabolism, we next performed untargeted high-resolution metabolomics^18^. Worms were fed *bcat-1* or control RNAi as adults and collected on day 5, the same timepoint as our transcriptomics analysis. Significantly altered features (*P*<0.05) were detected in *bcat-1(RNAi)* worms compared with control worms (Fig. 2a,b, Extended Data Table 2), and PLS-DA showed distinct separation of the groups (Fig. 2c).

**Figure 2:**
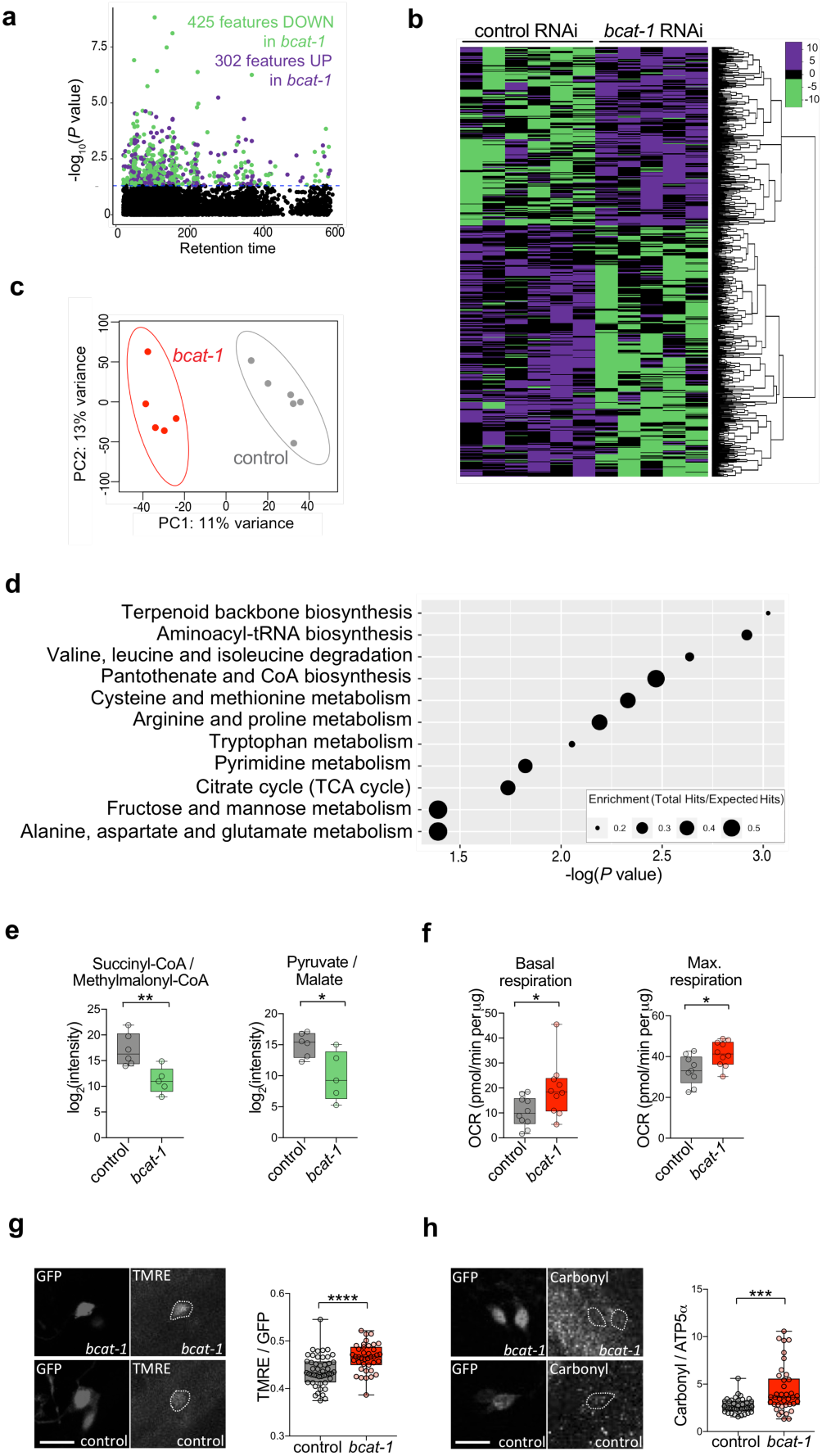
Depletion of TCA cycle metabolites coincides with increased mitochondrial activity in *bcat-1(RNAi)* worms. **a-e**, Neuronal RNAi-sensitive worms were collected for metabolomics on day 5 following adult-specific *bcat-1* knockdown. Manhattan plot **(a)** and heat-map **(b)** show features increased (purple) and decreased (green) (*P*<0.05) in *bcat-1(RNAi)* worms. Color values in **(b)** represent relative intensities of features that have been log2-transformed and centered, **c**, Partial least squares—discriminant analysis, **d**, Pathway analysis using mummichog algorithm, **e**, TCA cycle metabolites/precursors were decreased in *bcat-1(RNAi)* worms. n=6 independent collections for control, 5 for *bcat-1*. **f**, Mitochondrial respiration was increased on day 5 in *bcat-1(RNAi)* worms. n=10 wells totaling 102 worms for control basal, 9 wells totaling 92 worms for control max., 10 wells totaling 100 worms per *bcat-1* condition, **g**, Mitochondrial activity was increased on day 5 in CEP α-synuclein -expressing dopaminergic neurons with *bcat-1* knockdown. Scale bar, 10μm. n=15 worms totaling 47 CEPs for control, 11 worms totaling 39 CEPs for *bcat-1*. **h**, Protein carbonylation was increased on day 8 in CEP α-synucleinexpressing dopaminergic neurons with *bcat-1* knockdown. Scale bar, 10μm. n=10 worms totaling 37 CEPs for control, 10 worms totaling 38 CEPs for *bcat-1*. Two-tailed *t*-tests. \**P*<0.05, \*\**P*<0.01, \*\*\**P*<0.001, \*\*\*\**P*< 0.0001. Box-plots show minimum, 25th percentile, median, 75th percentile, maximum.

Several of the metabolic pathways that were significantly altered in *bcat-1(RNAi)* worms were related to BCAA metabolism, including *valine, leucine, and isoleucine degradation*, and *alanine, aspartate, and glutamate metabolism* (Fig. 2d). As expected, features putatively annotated as leucine/isoleucine and glutamate were increased or decreased, respectively, in *bcat-1(RNAi)* worms (Extended Data Fig. 4). Consistent with our transcriptomics analysis, *citrate cycle (TCA cycle)* was also a significantly altered metabolic pathway in *bcat-1(RNAi)* worms (Fig. 2d). However, while the expression of TCA cycle genes was largely increased (Fig. 1i), the levels of the annotated TCA cycle substrates and precursors pyruvate/malate and succinyl-CoA/methylmalonyl-CoA were significantly decreased (Fig. 2e). These findings suggest that *bcat-1* knockdown reduces steady-state levels of TCA cycle metabolites, while increasing expression of glycolysis and TCA cycle enzymes.

It is possible that low levels of TCA cycle metabolites leads to a reduction in mitochondrial respiration. Alternatively, low steady-state levels of metabolites may reflect more rapid TCA cycle turnover driven by more abundant components, resulting in higher levels of mitochondrial respiration. To distinguish between these possibilities, we measured mitochondrial activity in worms with *bcat-1* knockdown. Oxygen consumption rates were assayed on day 5, and both basal and maximal respiration rates were found to be increased in *bcat-1(RNAi)* worms (Fig. 2f). Similarly, using TMRE dye that accumulates only in active mitochondria, we found that dopaminergic neurons expressing α-synuclein had increased mitochondrial activity with *bcat-1* knockdown (Fig. 2g, Extended Data Fig. 5a). These data support the conclusion that *bcat-1* reduction leads to mitochondrial hyperactivity.

Since mitochondrial respiration produces reactive oxygen species, we next tested for evidence of oxidative damage. Protein carbonylation levels were measured in α-synuclein-expressing dopaminergic neurons with or without *bcat-1* knockdown at the day 8 endpoint. Indeed, high mitochondrial activity was correlated with elevated levels of carbonylated proteins (Fig. 2h, Extended Data Fig. 5b), suggesting that disruption of *bcat-1* dysregulates the cellular redox state such that neurons incur protein damage.

## Metformin reduces neurodegeneration and restores normal mitochondrial activity

While disease mechanisms in PD are thought to involve decreased mitochondrial function^19^, our findings instead suggest that elevated mitochondrial activity may be a causative factor in PD. Strategies to restore mitochondrial homeostasis may therefore be efficacious. We recently identified the type 2 diabetes medication metformin as the top-performing candidate in a high-throughput screen for drugs that improve motor function in *bcat-1(RNAi)* worms^3^. Among several known targets, metformin appears to act as a complex I inhibitor^20^, raising the possibility that reducing mitochondrial activity may constitute a surprising new treatment avenue for PD.

To investigate the potential neuroprotective action of metformin and its effects on mitochondria in the *bcat-1* PD worm model, we used the experimental paradigm established in our high-throughput drug screen^3^. Neuronal RNAi-sensitive worms were fed *bcat-1* or control RNAi as adults, and on day 4 transferred to plates seeded with heat-killed OP50 *E. coli* and 50 µM drug or vehicle (0.05% DMSO) (Fig. 3a). The use of heat-killed bacteria eliminates potential confounding effects of bacterial metabolism of the drugs, as has been documented for metformin^21^. In addition, drug intervention on day 4 mimics the current clinical landscape, in which PD diagnosis and treatment occurs well into the disease course^22^. On day 8, curling was quantified using our automated system^3^. Exposure to *bcat-1* RNAi until day 4, followed by vehicle treatment was sufficient to produce significant curling on day 8 (Fig. 3a). Using this paradigm, two PD medications currently prescribed for motor symptoms significantly reduced curling, whereas a PD drug indicated for cognitive symptoms had no effect (Fig. 3b), supporting the notion that curling reflects the motility defects of PD.

**Figure 3:**
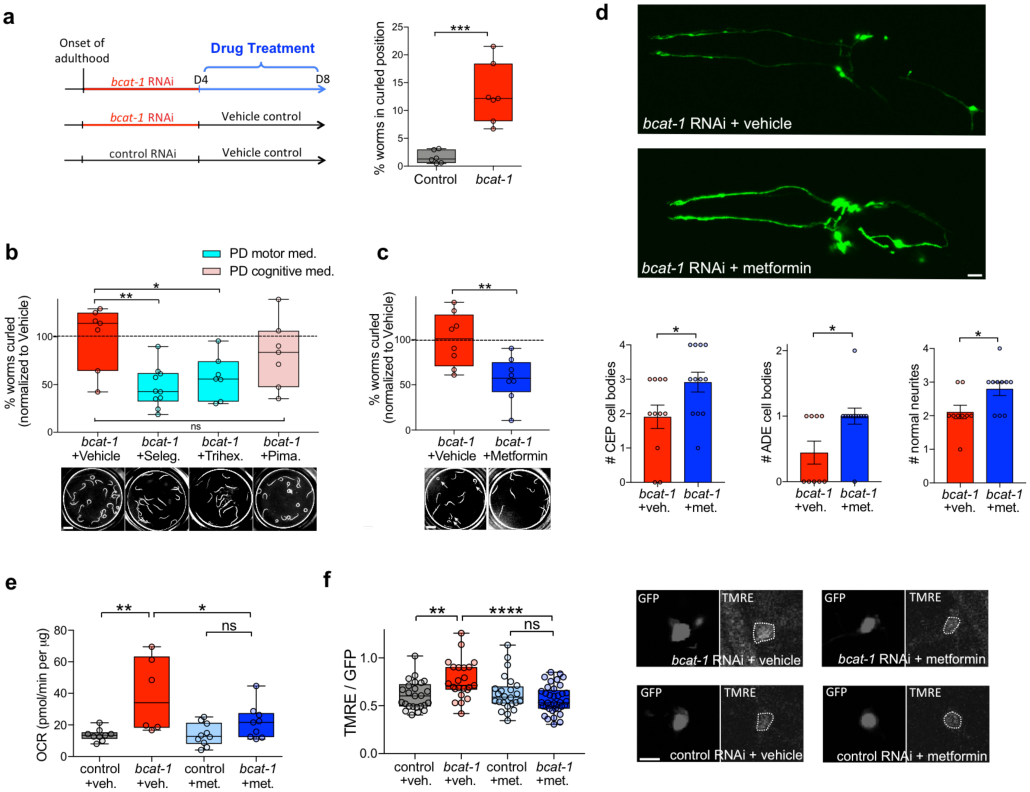
Metformin reduces neurodegeneration and restores normal mitochondrial activity in *bcat-1(RNAi)* worms. **a**, Experimental design for drug treatments. Neuronal RNAi-sensitive worms were fed *bcat-1* or control RNAi as adults until day 4, then transferred to heat-killed OP50 *E. coli* and 50 µM drug or vehicle. Vehicle-treated *bcat-1(RNAi)* worms curled on day 8, measured using our automated system. n=6 wells totaling 91 worms for control, 7 wells totaling 57 worms for *bcat-1*. Two-tailed *t*-test. **b**, PD medications prescribed for motor symptoms (selegiline (seleg.), trihexyphenidyl (trihex.)) reduced curling in *bcat-1(RNAi)* worms on day 8, whereas the antipsychotic pimavanserin (pima.) did not. Scale bar, 1mm. n=7 wells totaling 142 worms for *bcat-1*, 10 wells totaling 149 worms for selegiline, 7 wells totaling 86 worms for trihexyphenidyl, 7 wells totaling 103 worms for pimavanserin. One-way ANOVA with Dunnett’s post-hoc. **c**, Metformin reduced curling (arrows) on day 8 in *bcat-1(RNAi)* worms expressing α-synuclein in dopaminergic neurons. Scale bar, 1mm. n=8 wells totaling 65 worms for vehicle, 8 wells totaling 121 worms for metformin. Two-tailed *t*-test. **d**, Metformin reduced neurodegeneration of α-synuclein-expressing dopaminergic neurons with *bcat-1* knockdown on day 8. Scale bar, 10μm. n=11 for CEP vehicle, 12 each for CEP and ADE metformin, 9 each for ADE and neurites vehicle, 10 for neurites metformin. Two-tailed *t*-tests. Data are mean±s.e.m. **e**, Metformin reduced basal mitochondrial respiration in *bcat-1(RNAi)* worms on day 8. n=9 wells totaling 144 worms for control vehicle, 6 wells totaling 50 worms for *bcat-1* vehicle, 10 wells totaling 128 worms for control metformin, 9 wells totaling 113 worms for *bcat-1* metformin. Two-way ANOVA with Tukey’s post-hoc. **f**, Metformin reduced mitochondrial activity in α-synuclein-expressing CEP dopaminergic neurons with *bcat-1* knockdown on day 6. Scale bar, 5μm. n=8 worms totaling 25 CEPs for control vehicle, 7 worms totaling 23 CEPs for *bcat-1* vehicle, 9 worms totaling 24 CEPs for control metformin, 10 worms totaling 37 CEPs for *bcat-1* metformin. Two-way ANOVA with Tukey’s post-hoc. met., metformin. veh., vehicle. ns, not significant. \**P*<0.05, \*\**P*<0.01, \*\*\**P*<0.001, \*\*\*\**P*< 0.0001. Box-plots show minimum, 25th percentile, median, 75th percentile, maximum.

To determine if metformin rescues *bcat-1-*associated toxicity in worms with α-synuclein expression in dopaminergic neurons, we measured motor function and dopamine neuron degeneration. Worms treated with 50 µM metformin as described (Fig. 3a) showed reduced curling on day 8 (Fig. 3c). Remarkably, these worms also had significantly less neurodegeneration. On day 8, the number of dopaminergic cell bodies and neurites was increased in metformin-treated *bcat-1(RNAi)* worms compared with vehicle-treated worms (Fig. 3d). Neuroprotection was detected as early as day 6, after only two days of metformin treatment (from day 4 to day 6), in that dopaminergic neurons showed significantly improved neurite morphologies (Extended Data Fig. 6).

We next asked if metformin might exert its protective effects by suppressing *bcat-1(RNAi)-*driven mitochondrial hyperactivity. Indeed, we found on day 8 that metformin treatment had reduced both basal and maximal mitochondrial respiration down to levels of age-matched controls (Fig. 3e, Extended Data Fig. 7a). Similarly, using TMRE dye on day 6 (the latest timepoint permitted by the assay), metformin treatment was already sufficient to restore mitochondrial activity levels in α-synuclein-expressing dopaminergic neurons of *bcat-1(RNAi)* worms (Fig. 3f, Extended Data Fig. 7b). These data suggest that metformin is able to correct aberrant mitochondrial activity in the *bcat-1(RNAi)* worm model of PD.

## Conclusions

Here we demonstrate that defective BCAA metabolism recapitulates several major features of PD, including motor dysfunction and neurodegeneration. Despite an intriguing overlap of clinical and pathological features between BCAA metabolic disorders and PD^23^, a potential link remains largely unexplored. Maple syrup urine disease (MSUD) arises from mutations in BCKDHC subunits, and commonly presents with parkinsonian symptoms in adult patients^24^. Strikingly, neuronal loss in the substantia nigra has also been documented in MSUD^25^. Meta-analysis of genome-wide association studies revealed associations between PD and the BCAA pathway genes methylcrotonoyl-CoA carboxylase 1 (*MCCC1)* and branched-chain ketoacid dehydrogenase kinase (*BCKDK)*^26^, further implicating this pathway in disease risk. Consistent with this, metabolomic profiling of PD patient urine revealed that BCAAs were elevated relative to controls, and higher BCAA levels were correlated with greater disease severity^27^.

Our transcriptomics, metabolomics, and respiration experiment approaches led to the surprising finding that *bcat-1* reduction causes mitochondrial hyperactivity, consistent with reports from mice lacking the mitochondrial *BCAT* isoform^28^. These findings challenge the prevailing notion that mitochondrial function is solely decreased in PD, in particular due to complex I deficiency, which has been documented in peripheral tissues from PD patients^29^ and postmortem patient brain samples^30^. Critically, however, mitochondrial activity levels in PD patient brain prior to death remain unknown, and therefore it is possible that an increase in activity precedes the ultimate loss of function. Complex I inhibitors such as MPTP and rotenone are often used to model PD, and MPTP induces parkinsonism in humans^31^. However, MPTP produces acute neurodegeneration that does not model the progressive nature of PD^32^, and rotenone models suffer from issues of variability and subject mortality^33^. While these compounds may model endpoint processes in the disease, our study offers mitochondrial hyperactivity as a potential early pathological event in PD.

In support of the notion that hyperactive mitochondria may drive disease and therefore reducing activity levels back to normal may be protective, we found that metformin treatment restored normal mitochondrial activity and reduced both neurodegeneration and motor deficits in a model of PD driven by defective BCAA metabolism. Collectively, our work supports a metabolic origin of PD, and points to metformin as an existing FDA-approved drug with great promise for repurposing to PD.

## Methods

### *C. elegans* strains and maintenance

Worms were maintained at 20°C on standard nematode growth medium (NGM) plates or high growth medium (HG) plates, seeded with OP50 *Escherichia coli* or HT115 RNAi *Escherichia coli*, as indicated. The following strains were used in this study: wild-type worms of the N2 Bristol strain, neuron-only RNAi strain CQ511 [*sid-1(pk3321)*]; uIs69 [pCFJ90 (*myo-2p::mCherry, unc-119p::sid-1*)], and the following neuronal RNAi-sensitive strains, CF512 (*fem-1(hc17); fer-15(b26)*), CQ434 baIn11 [*dat-1p::α-synuclein; dat-1p::gfp*]; vIs69 [pCFJ90 (*myo-2p::mCherry + unc-119p::sid-1*)], and TU3311 uIs60 (*unc-119p::sid-1, unc-119p::yfp*).

### RNAi and drug treatments

Worms were synchronized from eggs by bleaching and plated onto HG plates seeded with OP50. At the L4 stage, worms were transferred to RNAi-seeded 100-mm NGM or HG plates containing carbenicillin and IPTG that were pre-induced with 0.1 M IPTG 1 hour prior to transfer. Worms were transferred to fresh RNAi plates every 2-3 days. In all experiments, control RNAi refers to empty vector pL4440 in HT115 *Escherichia coli*. For experiments with metformin or PD drugs, worms were transferred on day 4 to NGM plates seeded with 1 mL heat-killed OP50 bacteria and 50 µM metformin (Sigma), 50 µM selegiline (MedChem Express), 50 µM trihexyphenidyl (MedChem Express), 50 µM pimavanserin (MedChem Express), or vehicle (0.5% DMSO). OP50 was killed by incubation at 65°C for 30 min. The worms were transferred to fresh NGM plates with 1 mL heat-killed OP50 and drug or vehicle on day 6. For rapamycin experiments, rapamycin (LC Laboratories) was prepared in NGM media at a final concentration of 100 µM, and worms were maintained on rapamycin or vehicle plates from L4 to day 8 with transfer to fresh plates every 2-3 days.

### Curling quantification

Manual quantification of individual worms’ curling level was performed as previously described^2^, and defined as percentage of time spent curling. We recently developed an automated platform for high-throughput curling analysis^3^. Briefly, on the day of analysis, worms were washed twice rapidly with M9 buffer and dispensed into 96-well plates. 30-sec videos or a series of snapshots were obtained for individual wells containing 3-30 worms each. Our curling detection software quantified the # of worms in a curled position, divided by the total # of worms detected, and this was defined as percentage of worms in a curled position.

### Neurodegeneration assays

Animals were mounted on 2% agarose pads in M9 and sodium azide. For dopaminergic (*dat-1p::GFP*-labeled) neurons, scoring was done as previously described^2^. Briefly, images were obtained on a Nikon A1 confocal microscope at 40x magnification, and z stacks were processed in Nikon NIS elements software. CEP and ADE cell bodies were counted, as well as neurites projecting anteriorly from CEP cell bodies.

### Basal slowing behavioral assays

On day 5, CQ434 baIn11 [*dat-1p::α-synuclein; dat-1p::gfp*]; vIs69 [pCFJ90 (*myo-2p::mCherry + unc-119p::sid-1*)] worms were individually picked onto NGM plates seeded with OP50. After 2 min, the number of body bends was manually counted for 20 sec. Each worm was then picked onto an unseeded NGM plate. After 2 min, the number of body bends was manually counted for 20 sec. If a worm curled during a 20 sec measurement period, the 20 sec was restarted when the worm resumed normal movement. The assay could not be performed on day 6 due to excessive curling.

### Neuron isolation and transcriptomics

Adult neuron isolation was performed as described previously^14^. Briefly, TU3311 uIs60 (*unc-119p::sid-1, unc-119p::yfp*) worms were synchronized by bleaching and transferred to fresh *bcat-1* or control RNAi plates every 2-3 days starting at L4. Adult worms were separated from progeny every 2 days by sedimentation in M9 buffer. On day 5, worms were washed with M9 buffer and incubated for 6.5 min with lysis buffer (200 mM DTT, 0.25% SDS, 20 mM Hepes pH 8.0, 3% sucrose). The worms were washed with M9 buffer and incubated in 20 mg/ml pronase from *Streptomyces griseus* (Sigma) for 12-20 min with vigorous pipetting every 2-3 min. When worm bodies were dissociated and the solution became cloudy, 2% fetal bovine serum (Gibco) was added to stop the reaction, and the mixture was filtered through 5µm filters (Millipore). GFP-labeled neurons were sorted using a FACSVantage SE w/ DiVa (BD Biosciences; 488nm excitation for GFP detection), with age-matched N2 worms used a negative control for setting gates. RNA was extracted, DNase digested, and cleaned using Qiagen RNEasy Minelute columns as previously described^14,15^. RNA sequencing libraries were prepared using the SMARTer Stranded Total RNA kit v2-Pico input mammalian, as per manufacturer instructions. Libraries were pooled and sequenced on the Illumina HiSeq 2000 platform, and the Galaxy Workflow System was used to analyze the RNA sequencing data. Reads were mapped to the *C. elegans* genome (WS245) using RNA STAR, and mapped reads that overlap with gene features were counted using htseq-count (mode = union). Per-gene counts were used as input for the DESeq2 R package for differential gene expression analysis. Five total independent collections were obtained for each condition (*bcat-1* or control), and *bcat-1* curling was confirmed for each replicate on day 8. Upon publishing, raw sequencing reads will be made available at NCBI Bioproject: PRJNA599166.

### Gene Ontology and KEGG analyses

Lists of up- or downregulated genes in *bcat-1(RNAi)* neurons were analyzed using gProfiler^34^ with the following settings: *C. elegans* organism, only annotated genes, g:SCS threshold, user threshold .05, ENTREZGENE_ACC. REVIGO was used to cluster and plot GO terms with q-value < 0.05.

### High-resolution metabolomics

CF512 (*fem-1(hc17); fer-15(b26)*) worms were synchonized by bleaching and sterilized by incubation at 25°C from L2-L4. Worms were transferred to fresh *bcat-1* or control RNAi plates every 2-3 days starting at L4. Adult worms were separated from progeny every 2 days by sedimentation in M9 buffer. On day 5, worms were washed with M9 buffer and ∼500 worms were snap frozen in liquid nitrogen per replicate. In parallel for each sample, ∼500 worms were incubated in M9 for 1 hour to release gut bacteria, and then the starved worms were discarded and the bacteria were collected as a blank. Metabolites were extracted using acetonitrile (in a 2:1 ratio) which was added to all samples along with an internal standard. Bead-beating was used to disrupt the worm cuticle and improve extraction. Each sample was placed in the bead beater at speed 6.5 m/s for 30 seconds, allowed to equilibrate on ice for one minute, and placed in the beater for another 30 seconds, at the same speed. All processing was done either on ice or in a cold room. Non-targeted high-resolution mass spectrometry was run at the Clinical Biomarkers lab at Emory University using a HILIC column (positive mode) and C18 column (negative mode) using chromatographic methods previously described^35^. Mass spectral data was generated on an orbitrap mass spectrometer on full scan mode scanning for mass range 85 to 850 Da. Data were extracted using the R packages apLCMS^36^ and xMSanalyzer^37^. Features that were 1.5 times the intensity in the respective bacterial blank and were present in at least 9 of the 21 samples were further analyzed. Features were imputed if missing with half the value of the minimum abundance, normalized to protein content of the sample, and log2 transformed. All data processing, analysis and visualization was done in R version 3.6.0, using functions: gplots::heatmap2()^38^, ggplot2::ggplot()^39^, and mixOmics::plsda()^40^. Pathway analysis was done using the mummichog algorithm^41^ hosted on the metaboanlayst (www.metaboanalyst.ca) module “MS peaks to Pathway”^42^, using the *Caenorhabditis elegans* metabolic reference map available through KEGG. Five total independent collections were obtained for *bcat-1(RNAi)* and six independent collections were obtained for control RNAi. *bcat-1* curling was confirmed for each replicate on day 8.

### Oxygen consumption measurements

Whole-worm respiration was measured using a Seahorse XFe96 analyzer as previously described^43^. Briefly, worms were plated (5-25 worms per well) in 200 µL M9 buffer containing dilute heat-killed OP50 to prevent starvation. The same dilute heat-killed OP50 solution was used as a blank. Baseline respiration was measured, followed by injection of FCCP (10 µM, final concentration) to elicit maximal respiration, followed by sodium azide (40 mM, final concentration) to account for non-mitochondrial respiration. Measurements were normalized to protein content per well, determined by BCA assay.

### TMRE staining

NGM plates seeded with 1 mL heat-killed OP50 were spotted with tetramethylrhodamine ethyl ester (TMRE) to a final concentration of 30 µM and allowed to dry. Worms were transferred to TMRE plates, and the following day, animals were mounted on 2% agarose pads with levamisole. Images of dopaminergic cell bodies were obtained on a Nikon A1 confocal microscope at 100x magnification, and z stacks were processed in Nikon NIS elements software. For each cell, a region of interest was drawn around the cell body at the plane with the greatest GFP intensity. The corresponding TMRE intensity (TRITC channel) was normalized to the GFP intensity.

### Protein carbonylation staining

Staining was performed using the OxyBlot kit as previously described^44^. Briefly, worms were washed 3x with M9 and incubated in 1X DNPH for 30 min at room temperature. The reaction was blocked by adding 4x volume of neutralization buffer. Worms were washed 2x with ice-cold water, sandwiched between two poly-L-lysine coated slides, and incubated for 20 min in liquid nitrogen. The slide sandwiches were then freeze-cracked, the worms were fixed with methanol followed by acetone, and then blocked for one hour at room temperature with 1X PBS, 0.2% Gelatine, 0.25% Triton X-100. The primary antibody was applied at 1:100 dilution in blocking solution and incubated overnight at 4°C. As an internal control, anti-ATP5A antibody was co-incubated. Following four 10-min washed with 0.25% Triton X-100 in 1X PBS, secondary antibody was applied at 1:300 dilution in blocking solution and incubated overnight at 4°C. Images of dopaminergic cell bodies were obtained on a Nikon A1 confocal microscope at 60x magnification, and z stacks were processed in Nikon NIS elements software. For each cell, a region of interest was drawn around the cell body at the plane with the greatest GFP intensity. The corresponding anti-DNPH intensity (TRITC channel) was normalized to the anti-ATP5A intensity (Cy5 channel).

### Statistical analysis

For all comparisons between two groups, an unpaired two-tailed Student’s *t*-test was performed. For comparisons between multiple groups, One-Way ANOVA or Two-Way ANOVA (for 2 variables, i.e. RNAi treatment and +/- metformin) were performed with post-hoc testing as indicated. GraphPad Prism was used for all statistical analyses.

## Acknowledgments

We thank the *C. elegans* Genetics Center for strains (P40 OD010440), the Genomics Core Facility and Confocal Imaging Facility at Princeton University, Dr. Dean Jones and the Clinical Biomarkers Laboratory at Emory University, A. Tartici for help with data analysis, and the Murphy lab for discussion. C.T.M. is the Director of the Glenn Center for Aging Research at Princeton and an HHMI-Simons Faculty Scholar. D.E.M. was supported by Ruth L. Kirschstein NRSA (NIA F32AG062036) and further support was provided by NIH (NIEHS U2CES030163 and R01ES023839) to G.W.M., and NIH DP1 Pioneer Award to C.T.M. (NIGMS 5DP1GM119167) and The Glenn Foundation for Medical Research to C.T.M. (GMFR CNV1001899).

## Author Contributions

Conceptualization, D.E.M., R.K., and C.T.M.; Methodology, D.E.M., S.S., R.K., W.K., V.K., G.W.M., and C.T.M.; Investigation, D.E.M., S.S., R.K., W.K., and V.K.; Writing – Original Draft, D.E.M.; Writing – Review & Editing, D.E.M., R.K., V.K., G.W.M., and C.T.M.; Funding Acquisition, D.E.M., G.W.M., and C.T.M.

## Extended Data Figures and Tables

**Extended Data Fig. 1:**
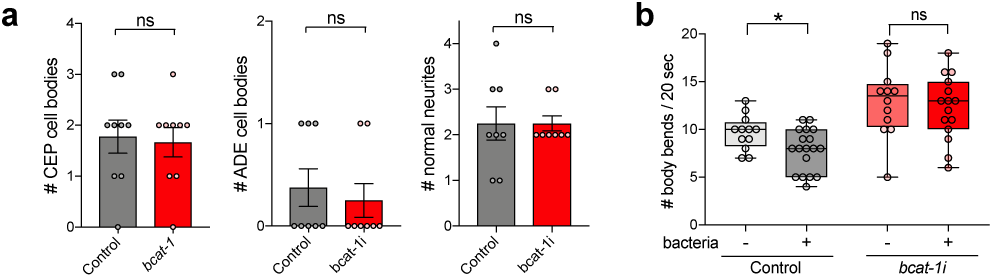
**a**, α-synuclein-expressing dopaminergic neurons with or without *bcat-1* knockdown are largely degenerated by day 8. n=9 each for control and *bcat-1* CEP, 8 each for control and *bcat-1* ADE and neurites. Two-tailed *t*-tests. Data are mean±s.e.m. **b**, While dopaminergic-dependent basal slowing behavior is still intact on day 5 in worms expressing α-synuclein in dopaminergic neurons, it is absent with additional knockdown of *bcat-1*. n=12 each for (-)bacteria, 18 for control (+)bacteria, 15 for *bcat-1* (+)bacteria. Two-tailed *t*-tests. ns, not significant. \**P*<0.05. Box-plots show minimum, 25th percentile, median, 75th percentile, maximum.

**Extended Data Fig. 2:**
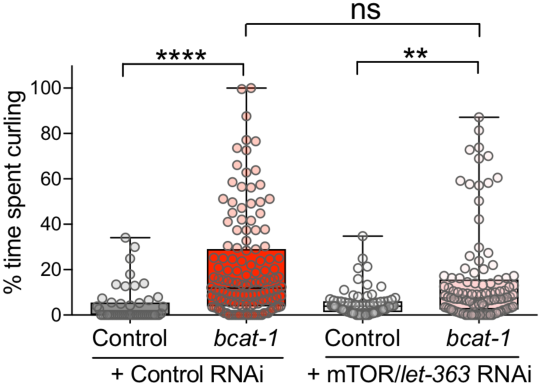
Curling is unaffected by mTOR/*let-363* knockdown in neuronal RNAi-sensitive worms with *bcat-1* knockdown on day 8. n=48 for control, 134 for *bcat-1* control, 72 for control *let-363*, 106 for *bcat-1 let-363*. Two-way ANOVA with Tukey’s post-hoc. ns, not significant. \*\**P*<0.01, \*\*\*\**P*< 0.0001. Box-plots show minimum, 25th percentile, median, 75th percentile, maximum.

**Extended Data Fig. 3:**
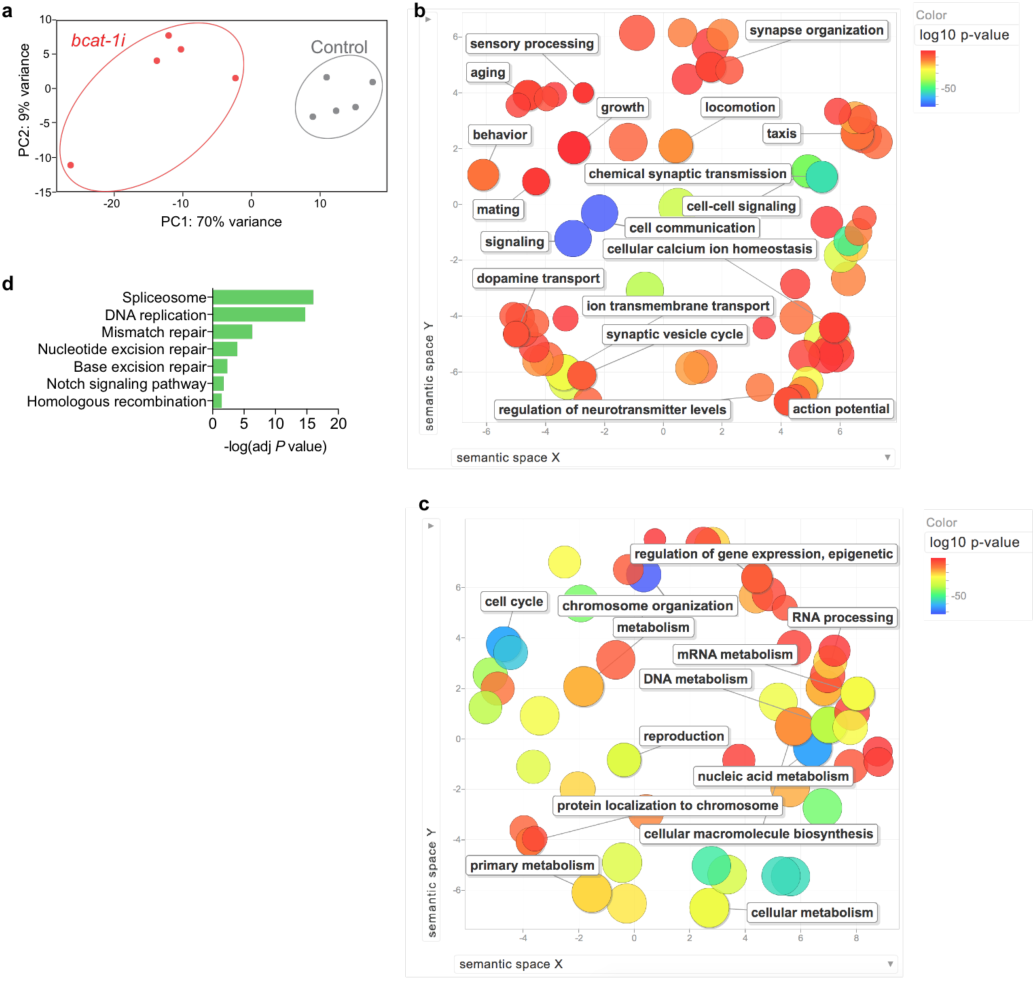
**a**, PCA plot of neuronal transcriptomes from worms with *bcat-1* knockdown and controls on day 5. **b-c**, Gene Ontology analysis of significantly upregulated (**b**) and downregulated (**c**) genes in *bcat-1(RNAi)* neurons. **d**, KEGG pathway analysis of significantly downregulated genes in *bcat-1(RNAi)* neurons. FDR<0.05. n=5 independent collections/group.

**Extended Data Fig. 4:**
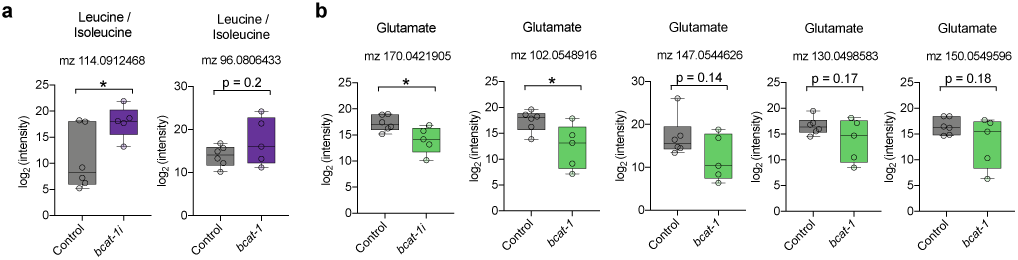
**a-b**, Levels of features annotated by mummichog as leucine/isoleucine (**a**) or glutamate (**b**) in *bcat-1(RNAi)* worms and controls. n=6 independent collections for control, 5 for *bcat-1.* Two-tailed *t*-tests. \**P*<0.05. Box-plots show minimum, 25th percentile, median, 75th percentile, maximum.

**Extended Data Fig. 5:**
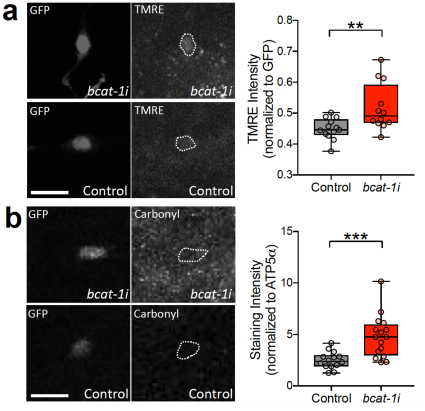
**a**, Mitochondrial activity was increased on day 5 in ADE α-synuclein-expressing dopaminergic neurons with *bcat-1* knockdown. Scale bar, 10µm. n=13 worms totaling 13 ADEs for control, 11 worms totaling 12 ADEs for *bcat-1*. **b**, Protein carbonylation was increased on day 8 in ADE α-synuclein-expressing dopaminergic neurons with *bcat-1* knockdown. Scale bar, 10µm. n=10 worms totaling 14 ADEs for control, 10 worms totaling 16 ADEs for *bcat-1*. Two-tailed *t*-tests. \*\**P*<0.01, \*\*\**P*<0.001. Box-plots show minimum, 25th percentile, median, 75th percentile, maximum.

**Extended Data Fig. 6:**
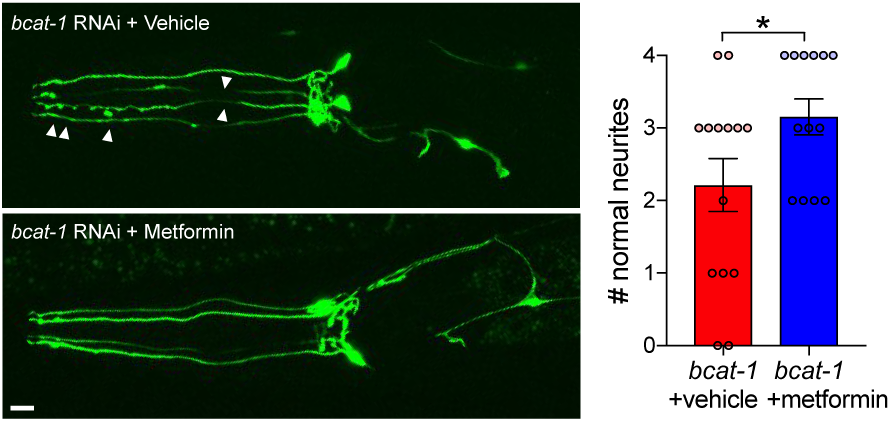
Metformin reduced degenerated morphologies (blebbing and fragmentation, arrowheads) in α-synuclein-expressing dopaminergic neurites with *bcat-1* knockdown on day 6, following only two days of treatment. Scale bar, 10µm. n=14 for vehicle, 13 for metformin. Two-tailed *t*-test. Data are mean±s.e.m.

**Extended Data Fig. 7:**
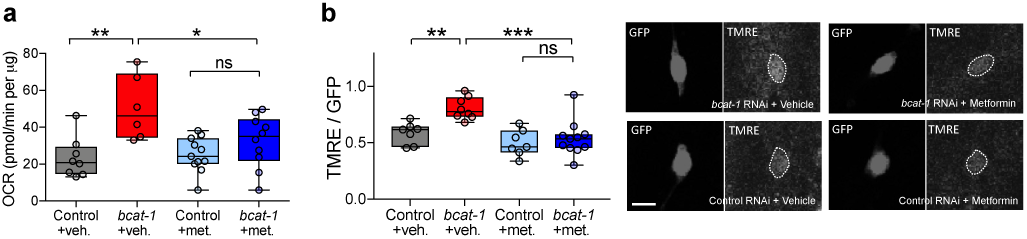
**a**, Metformin reduced maximal mitochondrial respiration in *bcat-1(RNAi)* worms on day 8. n=8 wells totaling 119 worms for control vehicle, 6 wells totaling 50 worms for *bcat-1* vehicle, 11 wells totaling 139 worms for control metformin, 10 wells totaling 123 worms for *bcat-1* metformin. Two-way ANOVA with Tukey’s post-hoc. **b**, Metformin reduced mitochondrial activity in α-synuclein-expressing ADE dopaminergic neurons with *bcat-1* knockdown on day 6. Scale bar, 5µm. n=7 worms totaling 7 ADEs for control vehicle, 7 worms totaling 8 ADEs for *bcat-1* vehicle, 7 worms totaling 7 ADEs for control metformin, 10 worms totaling 11 ADEs for *bcat-1* metformin. Two-way ANOVA with Tukey’s post-hoc. ns, not significant. \**P*<0.05, \*\**P*<0.01, \*\*\**P*<0.001. Box-plots show minimum, 25th percentile, median, 75th percentile, maximum.

**Extended Data Table 1: Differentially-expressed genes in *bcat-1(RNAi)* neurons.** Lists of all genes significantly up- or downregulated in *bcat-1(RNAi)* neurons (FDR<0.05), as well as subsets of each that were previously identified as neuronally-expressed^15^.

**Extended Data Table 2: Annotated features significant at p<0.05 from metabolomics analysis of *bcat-1(RNAi)* worms.** m.z, mass-to-charge ratio; retention time, retention time off of the column; KEGG.ID, KEGG ID determined by mummichog hosted on metaboanalyst; Annotation, Compound name from KEGG. (Level 5 confidence in annotation (Schymanksi 2014)); Adduct, Metabolite adduct annotated; Mass.Diff, Difference in mass of adduct detected and true mass of adduct.

